# Child specific activation in left auditory cortex predicts behavioral performance in inhibition tasks

**DOI:** 10.1101/2020.04.30.069906

**Authors:** Sam van Bijnen, Lauri Parkkonen, Tiina Parviainen

## Abstract

Sensory processing during development is important for the emerging cognitive skills underlying goal-directed behavior. Yet, it is not known how auditory processing in children is related to their cognitive functions. Here, we utilized combined magneto- and electroencephalographic (M/EEG) measurements to show that child-unique auditory cortical activity at ∼250 ms after auditory stimulation predicts the performance in inhibition tasks. While unaffected by task demands, the amplitude of the left-hemisphere activation pattern was significantly correlated with the variability of behavioral response time. Since this activation pattern is not present in adults, our results suggest divergent brain mechanisms in adults and children for consistent performance in auditory-based cognitive tasks. This difference can be explained as a shift in cortical resources for cognitive control from sensorimotor associations in the auditory cortex of children to top–down regulated control processes involving (pre)frontal and cingulate areas in adults.

## Introduction

The development of basic auditory circuits in the brain, and consequently efficient and versatile auditory behavior, relies on everyday aural experiences (Gordenet al., 2003; Tierney et al., 2015). Auditory sensory processing during development not only enables human communication and language learning, it also plays a role in cognitive and sensorimotor aspects of behavior (Kraus et al., 2012; Siegel et al., 2015). Indeed, the effect of auditory experience extends, for example, into attention and cognitive control processes that rely on auditory processing (Kraus and White-Schwoch, 2015). Presumably, an interaction between auditory, sensorimotor and cognitive processing govern the resulting phenotype of goal directed behavior (Kraus and White-Schwoch, 2015). Given the evident importance of auditory sensory development for cognitive skills, we have surprisingly limited understanding of how the typical development of cortical auditory processing is related to cognitive functions such as cognitive control.

Auditory evoked brain responses measured with electro- and magnetoencephalography (EEG/MEG) have been successfully used to study the development of the central auditory system (Paetau et al., 1995; Johnstone et al., 1996; Ponton et al., 2000; Ponton et al., 2002; Čeponienė et al., 2002; Wunderlich and Cone-Wesson, 2006) and they have been used as a marker for central auditory pathway plasticity (Sharma et al., 2002). Especially interesting from the perspective of auditory development is a prolonged activation pattern approximately 250ms after auditory stimulation, as it is typically reported in a wide age range of children but not in adults.

In adults, the resulting waveform from auditory stimulation is a combination of transient positive and negative deflections, which were defined by their order (P1-N1-P2-N2) or latency (e.g. N100) – and a lower letter “m” to indicate their MEG counterparts. In contrast, the most prominent responses in primary school children (∼6–12 years) are the P1(m) at around 100ms (Orekhova et al., 2013; Yoshimura et al., 2014) and a prolonged activation pattern at ∼250 ms (N2m/N250m) (Paetau et al., 1995; Ponton et al., 2000; Čeponienė et al., 2002; Parviainen et al., 2019). The development of the auditory neural activation is best characterized by a gradual dissociation of the earlier, more transient responses (P1/N1), and an attenuation of the later, prolonged, activity (N250) until it is no longer or barely present in adults (Ponton et al., 2000; Albrecht et al., 2000; Čeponienė et al., 2002; Takeshita et al., 2002; Wunderlich and Cone-Wesson, 2006). The right hemisphere seems to precede the left hemisphere in this developmental trajectory, suggesting faster maturation of the right-auditory cortex (Parviainen et al., 2019).

Developmental studies of human auditory processing have merely sketched the age-related changes in timing or strength of activation across the time-line of sensory activation. To go beyond the descriptive level, a fundamental question is how the development of activity in these time-windows (i.e. ∼100 and 250 ms.) is functionally meaningful for the development of cognitive functions. These two time-windows seem to represent functionally distinct processes. First, they are dissociated by their developmental trajectories (Parviainen et al., 2019). Second, activity in these time-windows show different refractory periods; whereas shortening the inter stimulus interval (ISI) attenuates the earlier response pattern, the later, prolonged activity is enhanced (or unaffected) (Takeshita et al., 2002; Karhu et al., 1997).

The child N1(m), emerging during early-mid childhood, seems to correspond relatively straightforward to the adult N1(m) (Čeponienė et al, 1998) and its role in auditory information processing is well known. In short, although the N1(m) primarily reflects sensory and perceptual processing, it is also affected by (selective) attention (Hilyard et al., 1973; Näätänen, 1982). In contrast, the later time-window (i.e. ∼200-300ms) shows remarkable differences between adults and children. Indeed, children show an auditory evoked response (i.e. N250m) that is reported even by passive stimulation, using different sound types, and under different attentional conditions (van Bijnen et al., 2019; Parviainen et al., 2019; Albrecht et al., 2000; Takeshita et al., 2002; Johnstone et al., 1996). This activity pattern is typically absent in adults. Instead, adults consistently show an evoked response in this time-window only in active tasks and it has been implicated in cognitive control in the cingulate cortex (Falkenstein et al., 1999; Nieuwenhuis et al., 2003; Huster et al., 2010). Accordingly, the general consensus of the time-line of auditory neural processing posits that the early transient peaks reflect stimulus-dependent processing, while beyond 100 ms the activation is thought to signify cognitive or evaluative aspects. Evidently, child-specific activation is likely to play a role in the functional development of auditory and related cognitive networks.

This prolonged activation pattern has been suggested to reflect increased automatization of information processing (Albrecht et al., 2000; Parviainen et al., 2011), possibly corresponding with the development of (neural) inhibition (Čeponienė et al., 2002) or the ability to control attention (Johnstone et al., 1996). However, direct correlational evidence comes only from language studies that have related weaker and/or contracted activity in this time window in typical developing children to a better performance on language tests (Parviainen et al. 2011; Hämäläinen et al., 2013). An empirical link between (the maturation of) this prolonged activity pattern and cognitive skills such as attention and inhibition has not been established.

Here, we utilized the excellent temporal accuracy of electrophysiological recordings and increased spatial sensitivity of combined MRI, MEG and EEG techniques to explore the behavioral significance of the child-unique activation at 250 ms. We focused on this activation pattern as it is most stable and prominent in the child brain in response to auditory stimulation, however, its functional significance remains elusive. We used comparisons between three variations of a simple auditory oddball paradigm (Fig. 1); a passive oddball task, a “detection” oddball task (press button for deviant tone) and an “inhibition” or Go/No-go task (press button for standard tone). Based on earlier findings we expected the child-specific auditory response to be present in children, but not adults, and we expected it to be present in both the active and passive (oddball & Go/No-go) tasks. We focused on (i) the effect of task on the amplitude of the auditory activation pattern in children and (ii) the relationship between this amplitude and behavioral performance measures of inhibition and/or attention (reaction time, response accuracy and intra-individual variability in reaction times).

**Figure 1.**
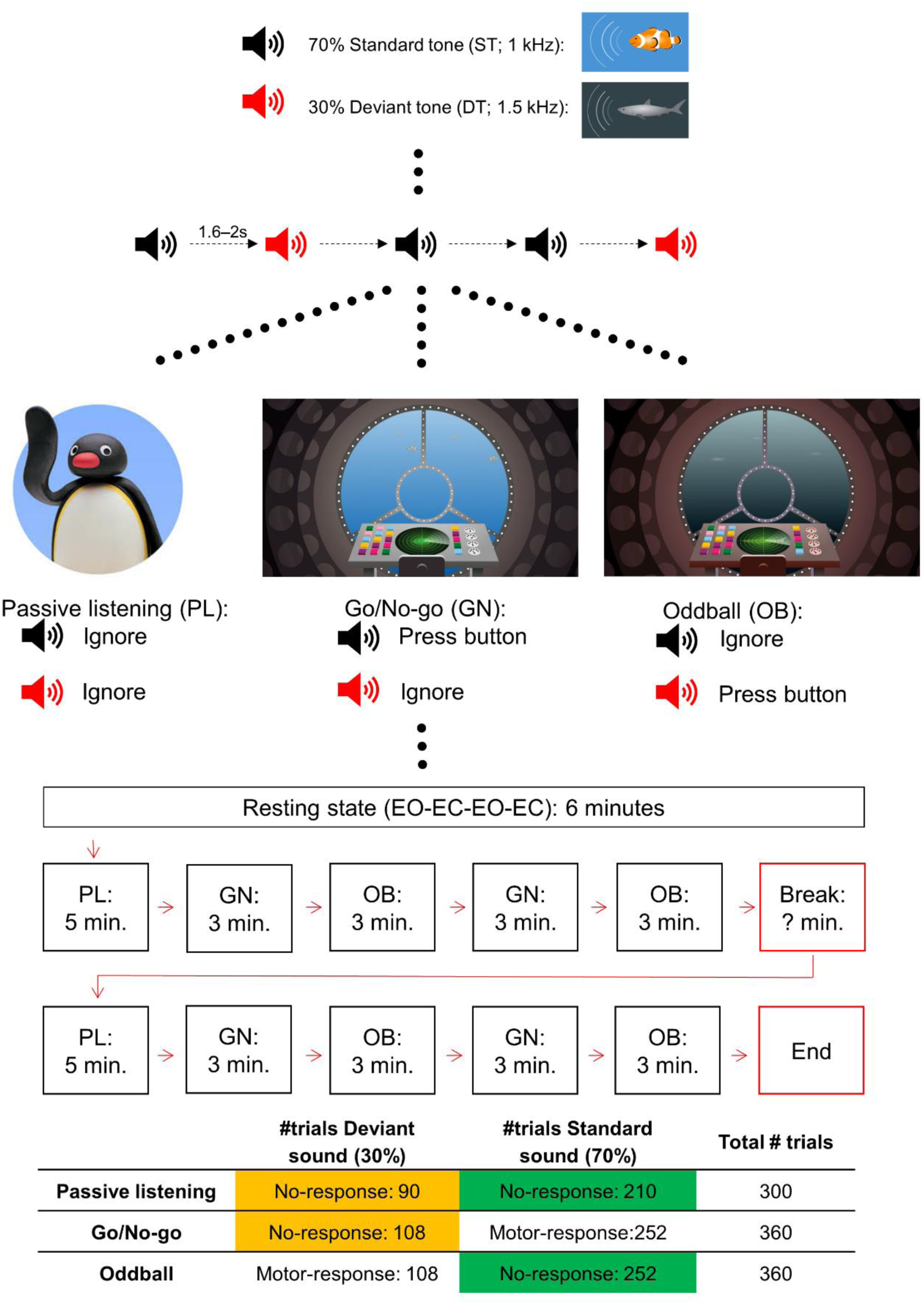
Experimental design and procedure. Statistical contrasts of interest marked in yellow/green (bottom table).

We combined M/EEG recordings and individual MRI’s to achieve maximal sensitivity to the spatiotemporal characteristics of maturation-specific activation patterns (Sharon et al., 2007). A combination of M/EEG is uniquely suitable to extract the separate components from the time-varying activation pattern evoked by auditory stimuli, and adding individuals MRI’s increases the accuracy of localizing the underlying cortical generators. Importantly for our purpose, while MEG is more sensitive to hemispheric differences, EEG provides a better account of deeper (e.g. cingulate cortex) and radial sources (Baillet, 2017; Gross, 2019).

## Materials and Methods

### Participants

Participants were Finnish speaking school children (6-14 years) recruited through schools and the National Registry of Finland, and Finnish speaking adults. None of the participants had neurological disorders or were on medication affecting the central nervous system. In total, 78 children and 16 adults participated in this study. Of the 78 children, eleven were excluded: one did not finish the experiment and one had too many errors in the MEG task (>50% errors in at least one block, see below), five had excessive head movements or magnetic interference during MEG/MRI measurements, two objected to go in the MRI scanner, and two showed structural abnormalities in their MRI. No adults were excluded. The data included in this study consisted of 67 children (mean age 10.2 years, SD: 1.4, range: 6‒14, 36 boys, 31 girls) and 16 adults (mean age 24.8, SD: 3.4, range: 20‒30, 3 men, 13 women). Children were recruited to cover mainly the ages between 8-12 years as previous studies indicated this age range is an important developmental period for our activation pattern of interest. All participants had normal hearing as tested with an audiometer. The study was approved by the Ethics Committee of the University of Jyväskylä. An informed consent was obtained from all children and their parents, and the adults in accordance with the Declaration of Helsinki. All participants received compensation for participation (movie ticket or gift card).

### Stimuli and Tasks

Auditory stimuli consisted of a 70-ms (10‒ms rise/fall time) sine wave tone with a frequency of either 1.0- (standard tone(ST); 70%) or 1.5-kHz (deviant tone(DT); 30%) at 65 dB SPL and were created with the Audacity software® (V2.3.3) (http://audacityteam.org/). A continuous stream of auditory stimuli was presented binaurally with an inter-stimulus interval varying between 1.6 and 2.0‒s. The stream always started with the standard tone, and two deviant tones were never presented in a row. The participants completed three tasks: a passive listening task (PL), an auditory Go/No-go (GN) and an auditory oddball task (OB). The stimuli were identical in all three tasks but the instructions on how to respond were different: subjects were asked to ignore the tones (PL), press a button to ST (GN), and press the button to DT (OB). The number of stimuli was different in the PL task compared to the GN and OB: 150 stimuli/block vs. 90 stimuli/block, respectively (Fig. 1).

The stimuli were embedded in a game. We created a visual environment resembling a submarine, where the captain gave instructions to the participants “inside” the submarine (Fig. 1). Visual stimuli were created by Studio Dennis Parren (www.dennisparren.com) and were there for the sole purpose of engaging the participants. All stimuli were controlled by PsychoPy (V3.2) (Peirce et al., 2019) running on a Linux desktop PC. Auditory stimuli were delivered to the subject through plastic tubes and earpieces using an MEG-compatible hi-fidelity sound system.

### Procedure

The experiment was conducted in a child-friendly environment in which the participants were asked to help science by studying the clownfish population. Before the start of the tasks, we measured resting-state activity with two times 1.5 minutes eyes open (EO) and eyes closed (EC). Subsequently, participants were instructed by a captain through movie clips on how to perform the three auditory tasks.

The first PL task started after the captain instructed the participant to ignore the tones while he would look for the clownfish. During this task, the participants watched the silent stop-motion animation series “Pingu”. After the first PL task, the captain explained that the submarine detects fish using sound (i.e., sonar) and that the captain needs help detecting them while he navigates the submarine. The participants were then told that the two tone-pips represented two types of fish (Fig. 1); the clownfish (ST) and the shark (DT). First, they were asked to detect the clownfish (GN task) by pressing a button (as quickly as possible) after the ST’s. Participants were also instructed to look in the middle of the window (Fig. 1) and focus on the sounds.

Twelve practice trials preceded the actual measurement to check whether the participants understood the task. Subsequently, in the OB task they were asked to detect the sharks by pressing a button whenever the DT was presented in order to protect the clownfish. Again, twelve practice trials were included to check whether the participants understood the task. Finally, two blocks of the GN task and OB task, each consisting of 90 trials (27 DT/63 ST), were completed alternately before the break. During the break, we offered participants a snack and drink and a possibility to stretch their legs. After the break, participants completed the same blocks again starting with the PL task followed by two blocks of alternating GN and OB tasks. The complete procedure is shown in Figure 1.

### M/EEG and MRI

The brain responses were recorded using a 306-channel MEG system and the integrated EEG system (Elekta Neuromag® TRIUX™, MEGIN Oy, Helsinki, Finland). M/EEG data were filtered to 0.1–330 Hz and sampled at 1000 Hz. EEG recordings were performed with a 32- channel cap and referenced online to an electrode on the right earlobe. Vertical and horizontal electrooculograms (EOG) were measured to capture eye movements and blinks for offline artifact suppression. EOG electrodes were placed directly below and above the right eye and on the outer canthi of each eye, and a common ground electrode was attached to the collarbone.

Five digitized head position indicator (HPI) coils were placed on the EEG cap to continuously monitor the head position in relation to the sensors of the MEG helmet. The EEG electrodes and HPI coils were digitized relative to three anatomic landmarks (nasion, left and right preauricular points) using the Polhemus Isotrak digital tracker system (Polhemus, Colchester, VT, United States). In addition, ∼150 distributed scalp points were digitized to aid in the co-registration with individual magnetic resonance images (MRIs).

T1- and T2-weighted 3D spin-echo MRI images were collected with a 1.5 T scanner (GoldSeal Signa HDxt, General Electric, Milwaukee, WI, USA) using a standard head coil and with the following parameters: TR/TE = 540/10 ms, flip angle = 90°, matrix size = 256 x 256, slice thickness = 1.2 mm, sagittal orientation.

### Behavioral assessment

Cognitive skills were tested on a separate visit. The behavioral tests included subtests of Wechsler Intelligence Scales for Children Third edition (Wechsler, 1991) or Wechsler Adult Intelligence Scale and the Stop Signal Task (SST) from the Cambridge Neuropsychological Automated Test Battery (CANTAB). Of the Wechsler Intelligence scale, the following subtests were administered: Similarities, Block Design, Digit Span, Coding and symbol search.

The similarities test is designed to assess verbal reasoning and the development of concepts. The block design subtest is designed to assess an individual’s ability to understand complex visual information. Digit span (backward/forward) is designed to measure verbal short-term memory and attention. The coding test is designed to measure speed of processing but is also affected by other cognitive abilities such as learning, short-term memory and concentration. Finally, the symbol search test is designed to measure processing speed but is also affected by other cognitive abilities such as visuomotor coordination and concentration.

In the SST, the participant must respond to an arrow stimulus by selecting one of two options depending on the direction in which the arrow points. The test consists of two parts: in the first part, the participant is first introduced to the test and told to press the left-hand button when they see a left-pointing arrow and the right-hand button when they see a right-pointing arrow. There is one block of 16 trials for the participant to practice this. In the second part, the participant is told to continue pressing the buttons when they see the arrows, but if they hear an auditory signal (a beep), they should withhold their response and not press the button. The task uses a staircase design for the stop signal delay (SSD), allowing the task to adapt to the performance of the participant, narrowing in on the 50% success rate for inhibition. The test is designed to measure response inhibition/impulse control.

### Data analysis

MEG data were first processed with the temporal signal space separation (tSSS) and movement compensation options, implemented in the MaxFilter™ program (version 3.0; MEGIN Oy, Helsinki, Finland), to suppress external interference and compensate for head movements (Taulu and Simola, 2006). The data were converted to the mean head position over the whole recording for each individual subject.

M/EEG data were analyzed using MNE-Python (version 0.17) (Gramfort et al., 2014; Gramfort et al, 2013). Continuous M/EEG recordings were low-pass filtered at 40 Hz, EEG data was re-referenced to the average over all EEG channels, and bad channels and data segments were identified and excluded. Epochs of –0.2 to 0.8 s relative to stimulus onset were then extracted and corrected for the baseline (–0.2 to 0s) offset. Epochs were rejected for incorrect responses and large MEG signals (> 4 pT/cm for gradiometers, > 5 pT for magnetometers). Independent component analysis (ICA) was applied to suppress ocular and cardiac artifacts separately for MEG and EEG (Hyvärinen and Oja, 2000). Next, *autoreject,* an automatic data-driven algorithm, was used on the EEG data to repair or exclude bad epochs. We followed procedure introduced by Jas and colleagues (2017). If the algorithm excluded more than 20% of the epochs, manual artifact rejection of the EEG epochs was used instead. Finally, the data were manually checked for obvious artifacts, and the six experimental conditions were averaged separately.

The cortical surface for the source model was constructed from the individual structural MRI with the Freesurfer software (RRID: SCR_001847, Martinos Center for Biomedical Imaging, http://freesurfer.net; Dale et al., 1999; Fischl et al., 1999; Fischl et al., 2001). The M/EEG source space was decimated at 4.9 mm spacing, resulting in ∼5000 current locations per hemisphere.

The MEG and EEG data were registered to the structural data with MNE coregistration using the fiducial landmark locations, digitized EEG electrode locations and the additional scalp point. A forward solution for the source space was constructed using three-layer BEMs. Conductivity values used for the intracranial tissue (brain, CSF), skull and scalp were set to 0.3, 0.006 and 0.3 for adults and 0.33, 0.0132 and 0.33 for children, respectively. The noise covariance matrix was calculated from the individual epochs 200-ms pre-stimulus baseline, using a cross validation method implemented in MNE. In order to combine data from the MEG gradiometers, MEG magnetometers and EEG electrodes into a single inverse solution, the forward solution matrix and data were whitened using the covariance matrix (Engemann and Gramfort, 2015).

The source currents were examined using a cortically-constrained, depth-weighted (*p* = 0.8) L2 minimum norm estimate (Hämäläinen and Ilmoniemi, 1994) with a loose orientation constraint (0.2). To determine the direction of the source currents, the source components normal to the cortical surface were extracted. The MNE solutions were constructed for each individual subject; source waveforms were computed as the mean value of the source element within region-of-interest (ROI) label 30 (transverse temporal gyrus) as defined by the Desikan-Killiany Atlas (Desikan et al., 2006). Amplitude values of the prolonged activity were calculated as an average over the 200-325ms time-window after stimulus presentation, which was determined by visual inspection of the grand averages (see Fig. 2). Only negative averages were included in the statistical analysis, as we assumed positive values would reflect cortical activity unrelated to our response of interest.

**Figure 2.**
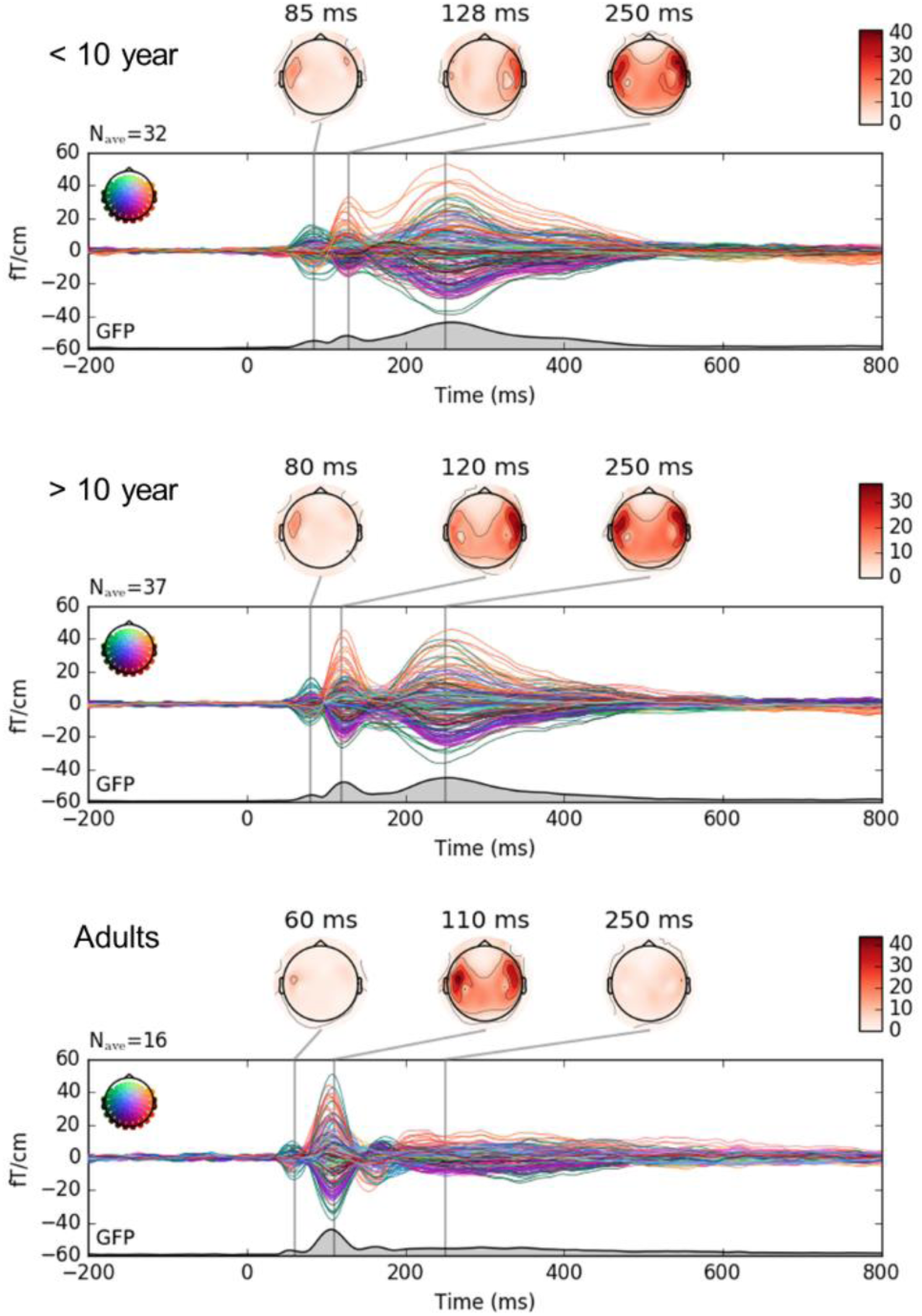
Developmental (age) differences in auditory brain responses to the passive listening (PL) standard tone (ST) as measured by the MEG gradiometers. Groups divided for illustration purposes between children younger than 10 (top), older than 10 (middle) and adults (bottom).

### Statistical analysis

As shown in Figure 1 (colored cells in bottom table) we designed the experiment to separately compare the effects of Oddball vs Passive (to focus on attention) and Go/No-go vs Passive (to focus on inhibition). We used the deviant tones (DT) for the comparison between Passive and Go/No-go (GN) and the standard tones (ST) for the comparison between Passive and Oddball (OB). Crucially, for these comparisons the stimuli (ST or DT), probability (30% or 70%) and motor response (None) were identical and the amount of trials close to equal.

A multiple linear regression analysis was performed to test for main effects of age, hemisphere and task. Subsequently, a related samples Wilcoxon Signed-ranks test was used to explore and describe the effects in more detail, as the brain response distributions were heavily skewed (non-negative values were excluded).

Partial correlations (controlling for age) were calculated for behavioral performance measures and the 2×2 (hemi x task) auditory brain responses. We included the following behavioral performance measurements: mean reaction time (RT), intra-individual coefficient of variation (ICV; calculated as SDRT/mean RT), response accuracy (ERR; calculated as square root of error %) from tasks completed inside the scanner, and the stop-signal reaction time (SSRT), which was completed outside the scanner during the behavioral assessment.

Linear regression analyses were performed with the behavioral performance measures as dependent variables. Age was entered first followed by the brain responses as independent variables. All variables in the linear regression model were selected based on the significant partial correlations. All statistical analyses were performed using SPSS statistics 25.

Finally, a bivariate correlation was used to check whether the brain responses were related to any of the subtests of the Wechsler Intelligence Scales for Children (i.e. digit span, symbol coding, symbol search, block design or similarities) to see if we had to control for possible intelligence effects.

## Results

### Descriptive statistics of cognitive skills and behavioral performance

Descriptive statistics of the children’s performance during the M/EEG experiment and their cognitive skills as per the behavioral assessment session are presented in Table 1.

**Table 1.** Mean, standard deviation (SD) and range of behavioral performance measures. Reaction times (RT), intra-individual coefficient of variation (ICV) and response accuracy (ERR) gathered from the Go/No-go task (GN) and the Oddball task (OB). Stop-signal reaction time (SSRT) was gathered from the stop-signal task during the behavioral assessment.

|  | Children |  |  | Adults |  |  |
| --- | --- | --- | --- | --- | --- | --- |
|  | Mean | SD | Range | Mean | SD | Range |
| Age (years) | 10.17 | 1.44 | 6-14 | 24.78 | 3.38 | 20-30 |
| **M/EEG experiment** |  |  |  |  |  |  |
| GN RT (ms) | 484.20 | 82.74 | 328-693 | 298.50 | 57.5 | 221-395 |
| GN ICV | 0.4 | 0.09 | 0.19-0.56 | 0.27 | 0.05 | 0.2-0.35 |
| GN ERR (√%) | 2.54 | 1 | 0.53-4.87 | 1.36 | 0.62 | 0-2.17 |
| OB RT (ms) | 480.67 | 82.03 | 234-728 | 303.69 | 53.85 | 214-420 |
| OB ICV | 0.38 | 0.11 | 0.18-0.82 | 0.21 | 0.04 | 0.14-0.32 |
| OB ERR (√%) | 1.78 | 0.85 | 0-3.87 | 0.7 | 0.36 | 0-1.18 |
| **Behavioral assessment** |  |  |  |  |  |  |
| SSRT (ms) | 205.94 | 56.20 | 87-351 | 140.81 | 32.62 | 80-198 |
| Digit span* | 10.55 | 2.65 | 5-17 | 18.06 | 2.89 | 14-26 |
| Symbol search* | 12 | 2.58 | 5-18 | 36.50 | 7.84 | 19-46 |
| Coding* | 10.88 | 2.98 | 4-19 | 81.56 | 9.35 | 66-103 |
| Block design* | 11.61 | 2.97 | 4-17 | 52.63 | 10.15 | 36-65 |
| Similarities* | 10.39 | 2.63 | 2-16 | 28.38 | 3.2 | 24-35 |

\* standardized score for children, raw score for adults

### Developmental trajectory of the auditory evoked responses

Figure 2 shows the measured neuromagnetic responses to the standard tones in the passive listening task at MEG sensor level (gradiometers). For visualization purposes, groups were separated by age (< 10 years old, > 10 years old and adults). The main activation in children is a prolonged activation pattern at around 250ms (N250m) in both hemispheres. The activation pattern of the older children in the earlier time window (∼100ms) starts to resemble that of the adults, but only in the right hemisphere. In contrast, the main activation in adults is evoked at around 100ms in both hemispheres.

### Prolonged activity at ∼250ms in auditory cortex is unique to the child activation pattern

Figure 3 shows the evoked activity between groups in the left and right transverse temporal gyrus with tasks overlaid. The maximum activation in children, emerging around 250ms appears to be similar across tasks including passive listening. In contrast, adults do not show a similar response in the auditory cortex in any of the tasks. Indeed, combined M/EEG source localization of the activation pattern show marked differences between adults and children (Fig. 4).

**Figure 3.**
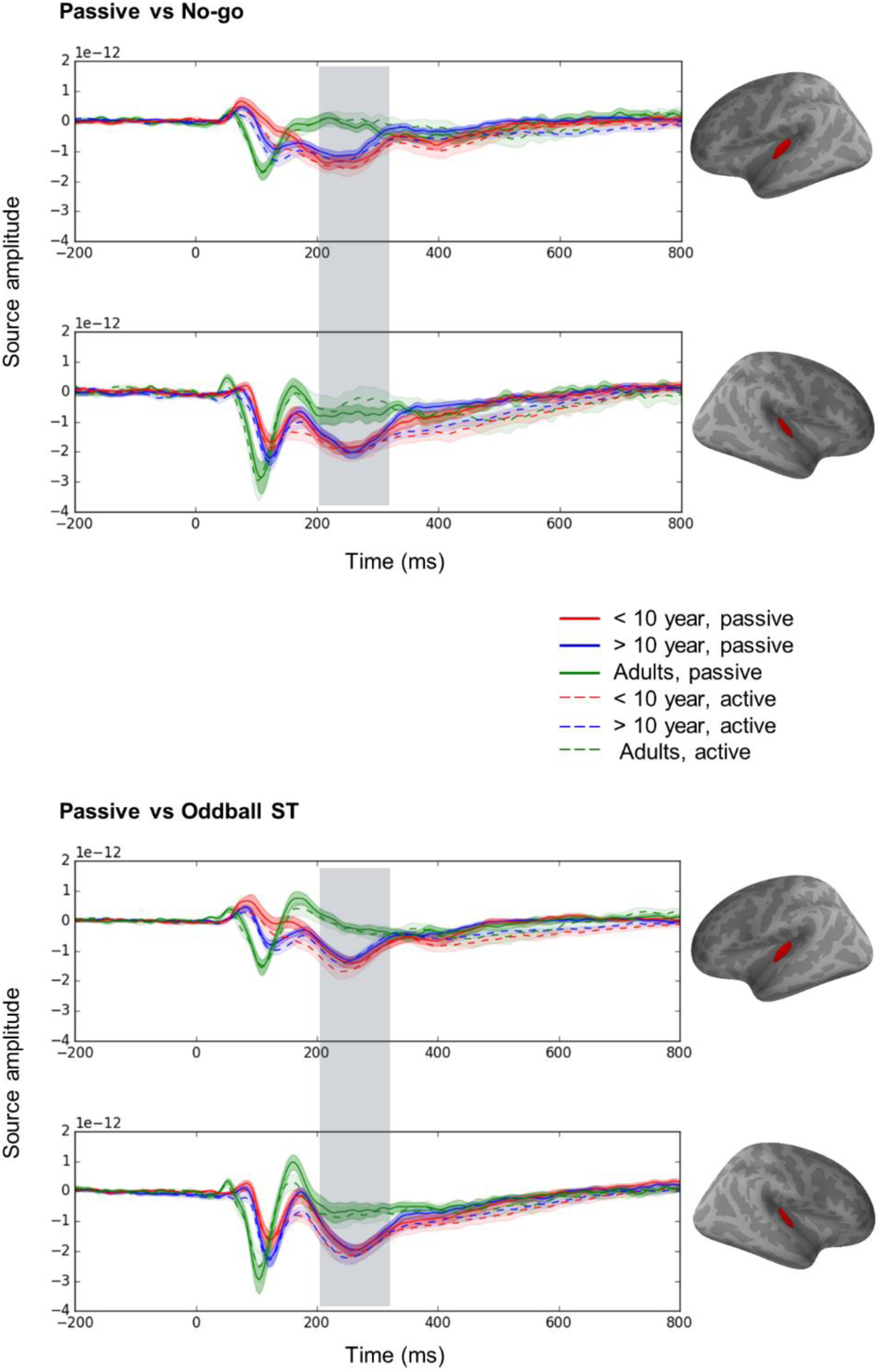
M/EEG combined Source estimates in the left and right transverse temporal gyrus (red area). Waveforms are an average of the entire area. Groups divided between < 10-year-old’s (red), > 10-year- old’s (blue) and adults (green). Top two figures depict the passive (solid lines) and attention (oddball standard tone) (dotted lines) waveforms in the left (top) and right (bottom) hemisphere. Bottom two figures depict the passive (solid lines) and inhibition (No-go deviant tone) (dotted lines) waveforms in the left (top) and right (bottom) hemisphere. Shaded areas around the waveform represent the standard error of the mean (SEM). Window is an approximation of the timepoints included in the calculation of the average.

**Figure 4.**
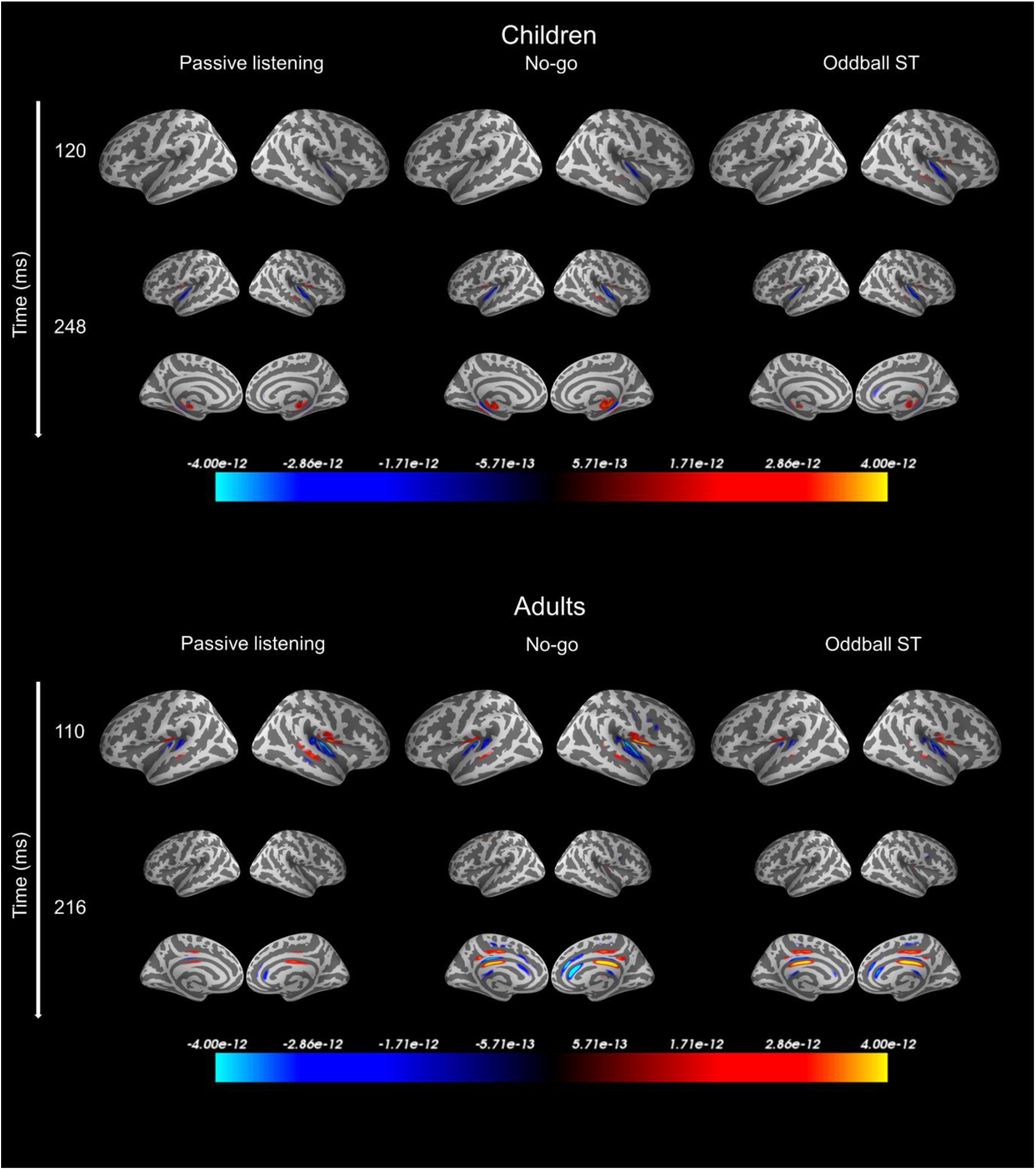
Grand average 3D visualization of the M/EEG combined source estimates for all children (top) and adults (bottom). 3D-plots presented for the two most prominent time-windows of activation in children (120ms and 248ms) and adults (110ms and 216ms). Conditions separated from left to right: Passive listening (standard tone(ST)), No-go (deviant tone) and Oddball (ST).

The source localization of the time-line of auditory neural processing tells us that the peaks in the child waveform are all localized in the temporal regions irrespective of task and time-window. In contrast, in adults the early peak at 100ms reflects activation in the temporal regions and the later activation ∼200-300ms reflects activation in the medial regions of the cerebral cortex (e.g., cingulate cortex). As the activation in children vs. adults in the 250- ms time-window reflect activation of different brain regions, their strength is not directly comparable. Moreover, the activation pattern at ∼250ms in the auditory cortex looks to be unique to the child brain (Fig. 3&4). Therefore, we did not directly contrast adults and children for this activation pattern. In the statistical analysis we focused on the strength of activation around ∼250ms after stimulus presentation in children’s transverse temporal gyrus. As per our experimental design, we discuss the PL vs GN and PL vs OB separately (see methods and Fig. 1).

### Passive vs Go/No-go

#### Right hemisphere shows generally stronger activation at ∼250ms independent of task

The multiple linear regression model, as shown in Table 2 revealed that hemisphere, but not age or task, was a significant predictor of the strength of activation in children. The Wilcoxon Signed-ranks test showed stronger activation in the right compared to the left hemisphere in both the PL and GN task. In the PL task the activation strength was 32% stronger in the right (*Mdn* = -15.18, IQR = [-8.85 – -21.81]) compared to the left hemisphere (*Mdn* = -10, IQR = [-5.35 – -13.6]), Z = -3.39, p = .001. Similarly, in the GN task the activation strength was 26% stronger in the right (*Mdn* = -16.82, IQR = [-9.57 – -24.18]) compared to the left (*Mdn* = - 11.29, IQR = [-5.6 – -17.58]) hemisphere, Z = -3.35, p = .001.

**Table 2.** Multiple linear regression analysis using hemisphere, task and age as predictors of the brain responses at ∼250ms.

| | *B* | *SE B* | Standardized beta | $\Delta R^2$ |
| --- | --- | --- | --- | --- |
| **Step 1** |  |  |  |  |
| **Constant** | -27.80 | 4.96 |  |  |
| **Hemisphere** | 4.82 | 1.19 | 0.25* | 0.08 |
| **Task** | -1.18 | 1.19 | -0.06 | ns |
| **Age** | 0.77 | 0.42 | 0.11 | ns |

Note: *B* = Unstandardized beta, *SE B* = standard error for the unstandardized beta, $\Delta R^2$ = $R^2$ change. \* $p < 0.05$ .

There was no significant effect of task on the activation strength. In general, the GN task showed non-significant stronger activation compared to the PL task. In the left hemisphere the activation in the GN task was 13% stronger (*Mdn* = -11.29, IQR = [-5.6 – -17.58]) compared to the PL task (*Mdn* = -10, IQR = [-5.35 – -13.6]), Z = -1.67, p = .095. In the right hemisphere, responses were 5% stronger in the GN task (*Mdn* = -16.82, IQR = [-9.57 – -24.18]) compared to the PL task (*Mdn* = -15.18, IQR = [-8.85 – -21.81]), Z = -0.82, p = .415.

Figure 5 shows the individual data points used the analysis as well as the average (line) and standard deviation (bar) for each condition.

**Figure 5.**
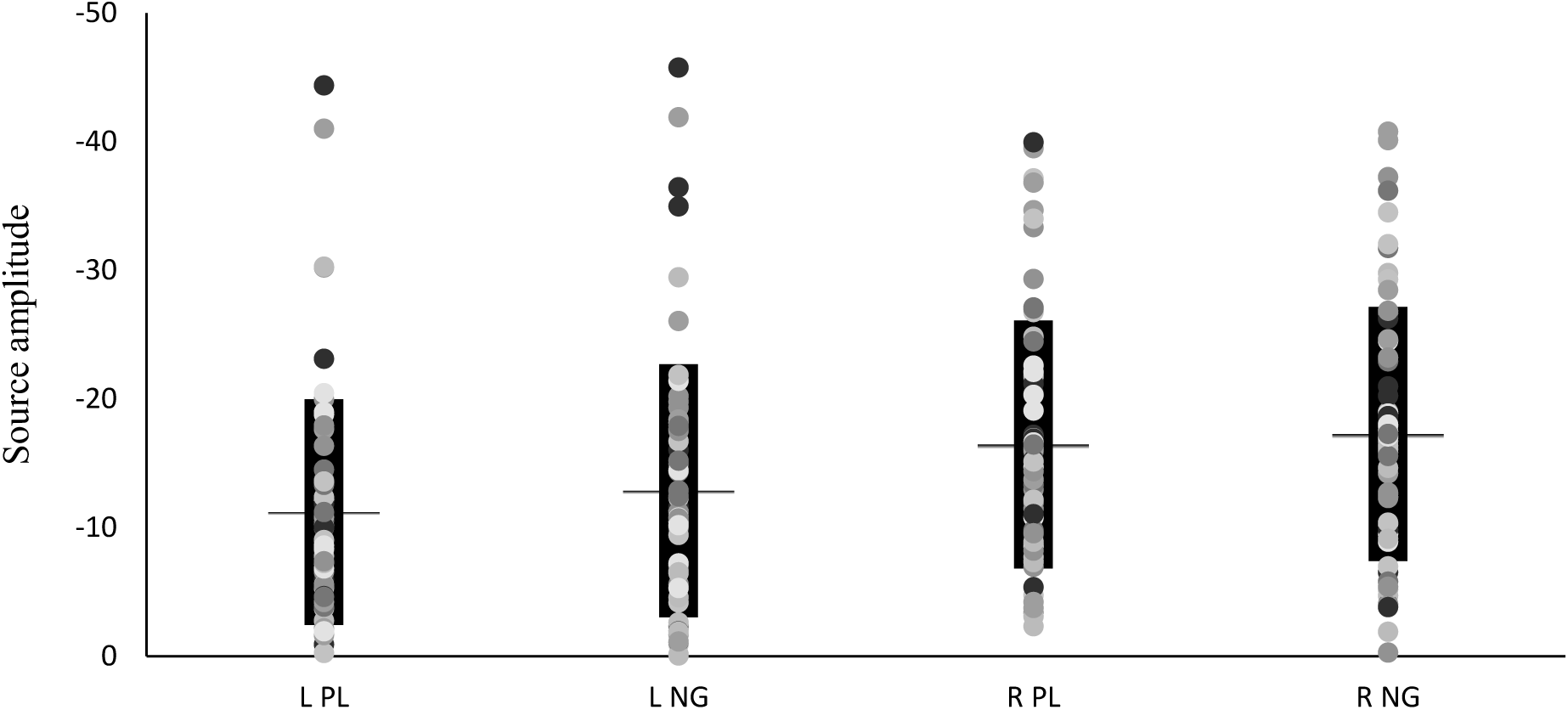
Individual data points (dots), average (horizontal line) and standard deviation (black bar) for the conditions: passive listening (PL) deviant tone and No-go (NG) deviant tone in the left (L) and right (R) hemisphere.

#### Left hemisphere auditory activity at ∼250ms predicts behavioral performance on inhibition tasks

A correlation analysis did not reveal any relationships between the brain responses and the subtests of the Wechsler Intelligence Scales for Children. As such, no control for general intelligence was added to the partial correlation analysis. Table 3 shows the result of a bootstrapped (10.000 samples) partial correlation (controlled for age) which revealed significant positive correlations between amplitudes in the left hemisphere (irrespective of task) and performance measures on both the Go/No-go (MEG inhibition task) and the SSRT (during behavioral assessment). Stronger left-hemisphere activation was related to lower intra-individual variability (ICV) in reaction times, lower error rate (ERR) and smaller stop-signal reaction times (SSRT).

**Table 3.** Bootstrapped (10.000 samples) partial correlation (controlled for age) between de brain responses and behavioral performance measures. Significant correlations marked in bold.

|  | RT | ICV | ERR | SSRT |
| --- | --- | --- | --- | --- |
| **L PL** | -0.024 | **0.479\*\*** | **0.314\*** | **0.331\*** |
| **R PL** | 0.157 | -0.033 | 0.037 | 0.162 |
| **L GN** | -0.019 | **0.467\*\*** | **0.343\*** | **0.292\*** |
| **R GN** | 0.035 | 0.077 | 0.036 | 0.231 |

*Note: RT = reaction time, ICV = intra-individual coefficient of variability, ERR = response accuracy, SSRT = stop signal reaction time. \* $p < 0.05$ , \*\* $p < 0.01$ .*

More specifically, in the PL task, a stronger left-hemisphere response amplitude was related to decreased ICV (r = .479, 95%CI = [.195 - .661], p = .000) and SSRT (r = .331, 95%CI = [.113 - .543], p = .02 and ERR (r = .314, 95%CI = [-.026 – .553], p = 0.028). Similarly, in the GN task, a stronger left-hemisphere response amplitude to the No-go tone was related to decreased ICV (r = .467, 95%CI = [.185 - .685], p = .001), decreased ERR (r = .343, 95%CI = [.022 - .587], p = 0.016), and decreased SSRT (r = .292, 95%CI = [.022 - .533], p = 0.041).

Subsequently, linear regressions were used to predict the performance measures using age and the selected brain responses. The brain responses to different tasks in the same hemisphere were highly correlated, and there was no significant effect of task, so we used the brain responses measured during the Go/No-go task. As shown in Table 4, the amplitude of the auditory response in the left hemisphere (to No-go tone) was a significant predictor of intra-individual variability of reaction time, error rate and stop-signal reaction time. Figure 6 shows the corresponding scatterplots.

**Figure 6.**
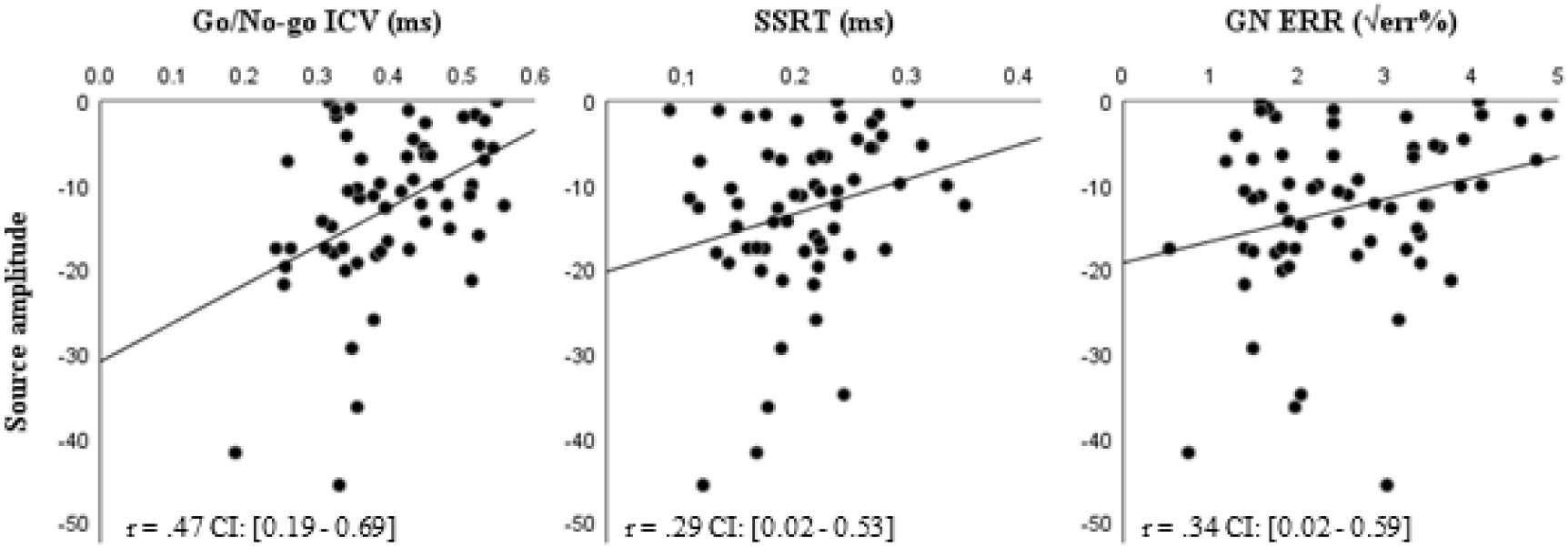
Scatterplots of the responses at ∼250ms to the No-go tone and the behavioral performance measures: intraindividual coefficient of variability (ICV; left), stop-signal reaction time (SSRT; middle), and response accuracy (right).

**Table 4.** Linear regression analysis using the behavioral performance measures as the dependent variable, age was entered first in the model, followed by the auditory responses in the left hemisphere to the No-go tone as the predictors.

| Performance measure | Step | Standardized Beta | $\Delta R^2$ |
| --- | --- | --- | --- |
| ICV | 1. Age | -0.248 | 0.036 |
|  | 2. Left auditory NG | 0.459 | **0.207\*\*** |
| ERR | 1. age | -0.319 | **0.078\*** |
|  | 2. Left auditory NG | 0.304 | **0.091\*** |
| SSRT | 1. age | -0.438 | **0.160\*\*** |
|  | 2. Left auditory NG | 0.295 | **0.086\*** |

Note: ICV = intra-individual coefficient of variability, ERR = response accuracy, SSRT = stop signal reaction time. \* $p < 0.05$ , \*\* $p < 0.01$ significance of $R^2$ change.

### Passive vs Oddball

#### Right hemisphere shows generally stronger activation at ∼250ms independent of task

Similar to the PL vs. GN comparison, the multiple linear regression model revealed that hemisphere, but not age or task, was a significant predictor of the strength of activation (see Table 5). The Wilcoxon Signed-ranks test showed significant stronger activation in the right compared to the left hemisphere in both the PL and OB task. The hemisphere effect was similar between tasks, with activation strength 29% stronger in the right (*Mdn =* -15.19, IQR = [-8.63 – -21.76]) compared to the left hemisphere (*Mdn* = -10.15, IQR = [-6.04 – -15.67]) in the PL task, Z = -3.329, p = .001, and 31% stronger in the right (*Mdn* = -18.27, IQR = [-10.4 – -22.56]) compared to the left hemisphere (*Mdn* = -10.82, IQR = [-6.8 – -16.11]) in the OB task, Z = - 4.24, p = .000.

**Table 5.**
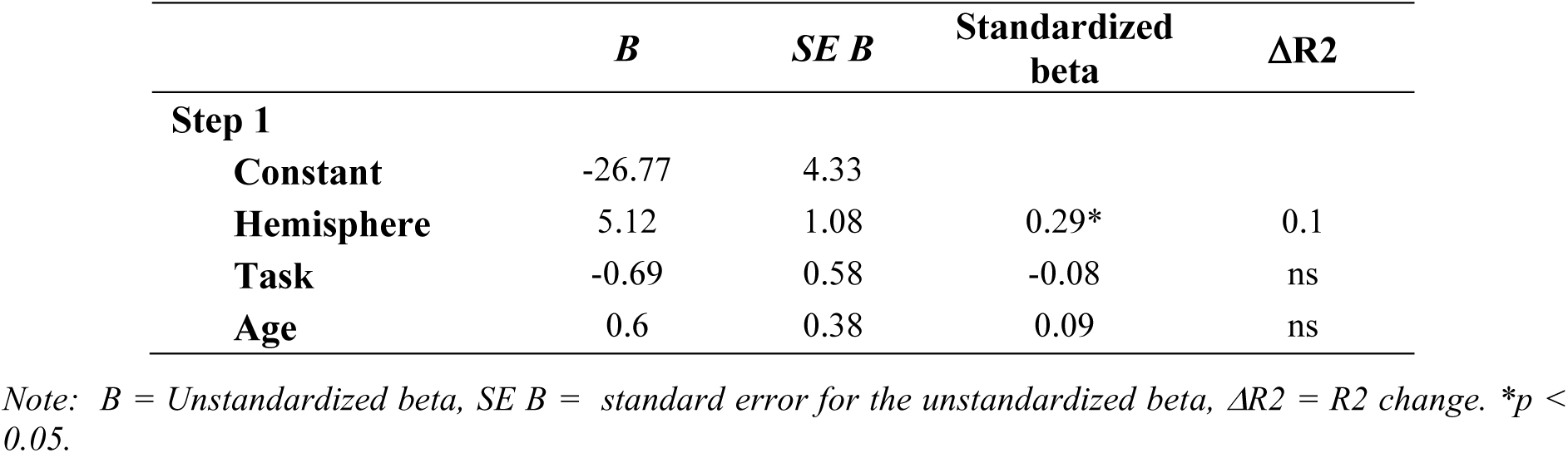
Multiple linear regression analysis using hemisphere, task and age as predictors of the brain responses at ∼250ms.

| | *B* | *SE B* | Standardized<br>beta | $\Delta R^2$ |
| --- | --- | --- | --- | --- |
| **Step 1** |  |  |  |  |
| **Constant** | -26.77 | 4.33 |  |  |
| **Hemisphere** | 5.12 | 1.08 | 0.29* | 0.1 |
| **Task** | -0.69 | 0.58 | -0.08 | ns |
| **Age** | 0.6 | 0.38 | 0.09 | ns |

Note: *B* = Unstandardized beta, *SE B* = standard error for the unstandardized beta, $\Delta R^2$ = $R^2$ change. \* $p < 0.05$ .

There was no significant effect of task. In the left hemisphere, activation strength was 8% stronger in the OB task (*Mdn* = -10.82, IQR = [-6.8 – -16.11]) compared to the PL task (*Mdn* = -10.15, IQR = [-6.04 – -15.67]), Z = -1.56, p = 0.119. In the right hemisphere, activation strength was 11% stronger in the OB (*Mdn =* -18.27, IQR = [-10.4 – -22.56]) compared to the PL task (*Mdn =* -15.19, IQR = [-8.63 – -21.76]), Z = -3.42, p = .001.

Figure 7 shows the individual data points used the analysis as well as the average (line) and standard deviation (bar) for each condition.

**Figure 7.**
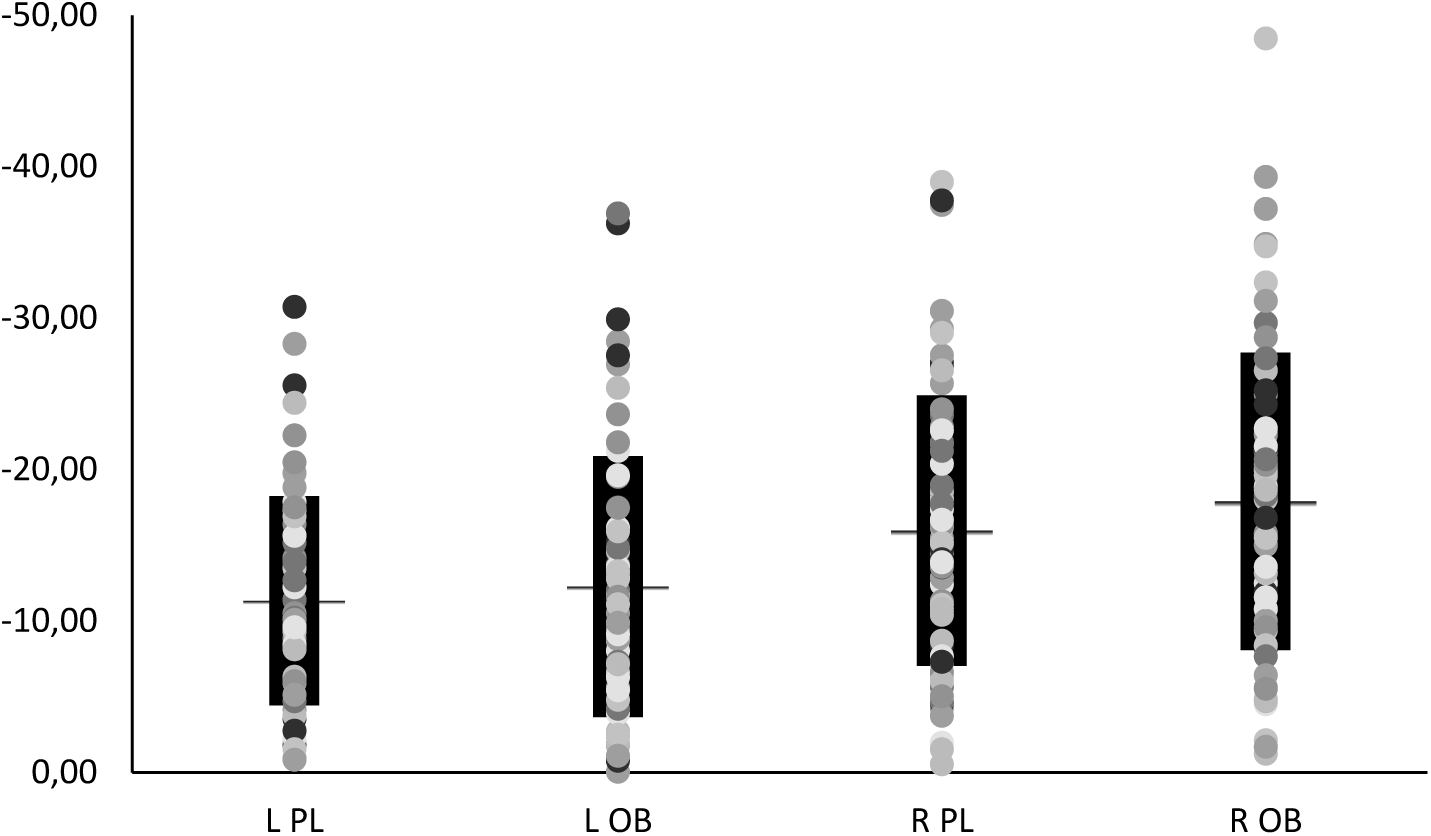
Individual data points (dots), average (horizontal line) and standard deviation (black bar) for the conditions: passive listening (PL) standard tone and oddball (OB) standard tone in the left (L) and right (R) hemisphere.

#### Left hemisphere auditory activity at ∼250ms predicts stop-signal reaction time

A correlation analysis showed no consistent relationships between the brain responses and the subtests of the Wechsler Intelligence Scales for Children; PL standard tone in the right hemisphere correlated with symbol search score (r = .261, p = .03) and the PL standard tone in the left hemisphere correlated with coding score (r = -.259, p = .04). No control for general intelligence was added to the partial correlation analysis. Table 6 shows the result of a bootstrapped (10.000 samples) partial correlation (controlled for age) revealed significant positive correlations between amplitudes in left hemisphere during the OB task and SSRT. Stronger activation in the left hemisphere during the OB task were related to smaller SSRT’s (r = 0.355, 95%CI = [0.142 – 0.560], p = 0.008).

**Table 6.** Bootstrapped (10.000 samples) partial correlation (controlled for age) between de brain responses and behavioral performance measures. Significant correlations marked in bold.

|  | RT | ICV | ERR | SSRT |
| --- | --- | --- | --- | --- |
| **L PL** | -0.153 | 0.252 | 0.194 | 0.251 |
| **R PL** | 0.087 | 0.042 | 0.025 | 0.224 |
| **L OB** | 0.033 | 0.234 | 0.230 | **0.355\*\*** |
| **R OB** | 0.143 | 0.086 | 0.028 | 0.238 |

*Note: RT = reaction time, ICV = intra-individual coefficient of variability, ERR = response accuracy, SSRT = stop signal reaction time. \*p < 0.05, \*\*p < 0.01.*

As shown in Table 7, the linear regression model revealed that the strength of the auditory response in the oddball task was a significant predictor of the SSRT (p = 0.019).

**Table 7.**
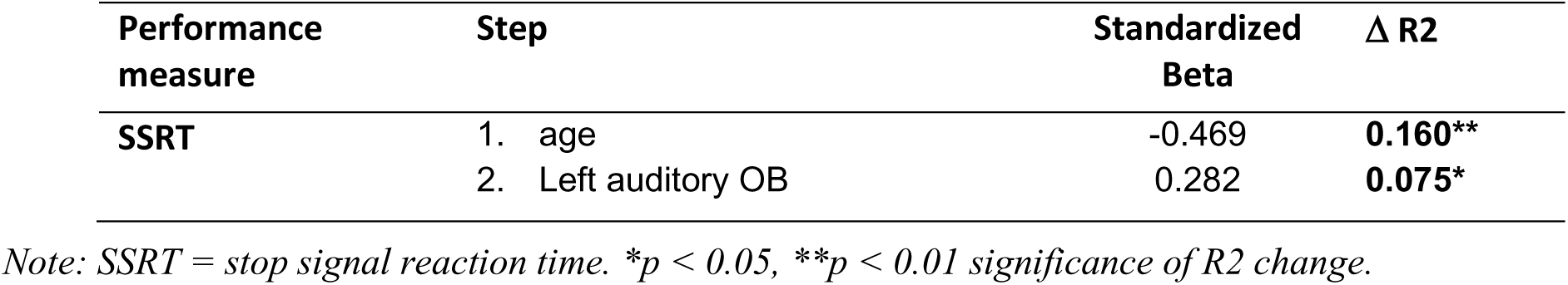
Linear regression analysis using the behavioral performance measures as the dependent variable, age was entered first in the model, followed by the auditory responses in the left hemisphere to the No-go tone as the predictors.

| Performance measure | Step | Standardized Beta | $\Delta R^2$ |
| --- | --- | --- | --- |
| **SSRT** | 1. age | -0.469 | **0.160\*\*** |
|  | 2. Left auditory OB | 0.282 | **0.075\*** |

*Note: SSRT = stop signal reaction time. \*p < 0.05, \*\*p < 0.01 significance of R2 change.*

## Discussion

We assessed the developmental trajectory and especially the functional significance of a robust activation pattern at ∼250ms (N250m) in children. The advanced source modelling of MEEG data confirmed that this activation pattern is uniquely prominent in the child brain; adults show an activation pattern in this time-window only in the active tasks and in different brain regions than children (i.e., medial regions of the cerebral cortex vs. auditory cortex, respectively). Age of the children did not seem to affect the strength of activation in this time window. This suggests a non-linear decrease during development of this auditory activation pattern with age, as it is clearly absent in adults. This was also indicated by previous studies with a wider age range than the present study, which found an initial increase in activation strength until children reached the age of 11, after which a gradual decrease was reported (Ponton et al., 2000; Ponton et al., 2002). Importantly, the strength of activation in children in the 250-ms time-window was unaffected by task demands, but the strength of activation in the left hemisphere was associated with superior performance on inhibition tasks and measures of cognitive control.

Our results confirm that (i) the N250m does not reflect a delayed adult N1m, nor does it correspond to the activation around 200ms in adults in active tasks (N2m) but instead is a developmentally specific auditory evoked brain response (Albrecht et al., 2000; Ponton et al., 2000, 2002; Takeshita et al., 2002; Parviainen et al., 2011; Parviainen et al., 2019) and that (ii) this prolonged activity pattern reflects general and automatic, circuit-level response characteristics in auditory areas of the child brain (Parviainen et al., 2019). Our findings bring important novel understanding of the functional significance of the child-specific activation pattern for the developing skills in cognitive control. It seems that engagement of the basic auditory cortex circuitry in the left hemisphere makes children more efficient in cognitive control as measured by the ICV.

The strength of the prolonged activation in the left, but not right, hemisphere was most consistently associated with performance on inhibition tasks. Left-hemisphere response strength explained 20.7%, 9.1% and 8.6% of unique variance of the ICV, response accuracy and SSRT respectively. We focus on the ICV and the prolonged activation during No-go trials, as the other results are likely different, less sensitive, measures of the same effect (i.e., one underlying effect is the most parsimonious explanation of our results).

The ICV reflects temporal variation in cognitive performance and it has been extensively studied in attention-deficit/hyperactivity disorder (ADHD) (de Zeeuw et al., 2008; van Belle et al., 2015). Intrasubject variability has long ago been put forward as an endophenotype of ADHD, the characteristic lapses of intention and attention in ADHD are thought to be a result of deficits in temporal processing that result in higher intrasubject intertrial variability (Castellanos and Tannock, 2002). Importantly, the auditory cortex coordinates activity with intricate timing. Indeed, the evoked responses reflect the auditory system’s ability to consistently respond with the same timing to each stimulus presentation. The behavioral importance of temporal processes is further supported by our and other studies’ finding that ICV, while unrelated to reaction time, is a much better predictor of inhibitory success (r = .79) than traditional measures of reaction time (r = .22) (Bellegrove et al., 2004; de Zeeuw et al., 2008, van Belle et al., 2015). Combined, these results suggest that ICV is an important measure of cognitive control that possibly relies on the auditory cortex’s (in auditory tasks) ability to consistently respond to the presented stimulus.

Our results indicate that the brain mechanisms that, in auditory based tasks, help achieve a consistent performance is remarkably different between children and adults. Most notably, the No-go activation in the 200-325ms time-window shows clear differences: whereas the adult major activation peak was localized to the medial regions of the cerebral cortex (e.g. cingulate cortex), children’s strongest activation pattern was located in the auditory cortex. Importantly, our findings are in line with earlier fMRI study’s and M/EEG studies in adults that emphasize the importance of both the 200-300 time-window and the cingulate cortex in inhibition and cognitive control (Nieuwenhuis et al., 2003; Huster et al., 2010; Falkenstein et al., 1999; Smith et al., 2007; Botvinick et al., 2004; Chambers et al., 2009). In contrast to the mature brain, our data show that children rely strongly on activation in the auditory cortex during the 200-300 time-window and although also evoked without task demands, it contributes to task performance.

Our results further suggest that the activity pattern during auditory inhibition tasks (e.g., Go/No-go or SST) in children and adults are qualitatively different. Consequently, due to divergent cortical origins of the signals, it is not informative to compare amplitude measures between adults and children in this time-window. This is relevant especially for EEG studies with limited spatial sensitivity; electrical potentials originating in the auditory cortices summate at the vertex, generating one maximum on the head surface (Hari and Puce, 2017). Consequently, even though the main current source underlying the measured signal is different between adults and children, typical EEG-ERP analysis will have limited capacity to reveal this difference, and may also erroneously transfer spatial differences into amplitude effects. Taken together, these results suggest that in order to move forward in understanding the neurodevelopmental underpinnings of improvement in cognitive skills (or problems therein), we need to adopt a more comprehensive approach in analysis incorporating both temporal and spatial characteristics of activation.

Our claim that children and adults employ different neural mechanisms to achieve a consistent performance in a cognitive control task is in line with previous fMRI studies. In adults, both reduced response variability and improved top-down cognitive control have been directly related to greater anterior cingulate gyrus (ACG) activity (Bellgrove et al., 2004; van Belle et al., 2015) and focal damage to the frontal lobes impairs the stability of cognitive performance (Stuss et al., 2003). In one fMRI study, younger subjects (7-15 years) showed differences from older subjects (15-24 years) in the relationship between dorsal ACG activity and response variability: in older children increased dorsal ACG activity was related to a reduction in response variability, whereas in the younger group dorsal ACG activity did not relate to this measure of cognitive control (van Belle et al., 2015). Intriguingly, Simmonds and colleagues (2007) reported that, in typically developing children (8-12 years), instead of cingulate activity, lower variability was associated with activation in the rostral supplementary motor area (pre-SMA) in a Go/No-go task.

The exact neurobiological underpinnings that underlie this difference between adults and children are unclear and should be the subject of further investigation. Our results together with earlier findings indicate a shift from sensorimotor associations in the child brain to more emphasis on cognitive control networks in the adult brain that underlie successful performance in inhibition tasks. In the present study, the strength of the prolonged activation in children showed a positive correlation with performance consistency, and thus seems to aid cognitive control in children. Importantly, we suggest a causal relationship based on the knowledge that this is an obligatory response to auditory stimulation, as it was similarly invoked without task demands. Thus, arguably, this child-specific sensory activation pattern enables flexible processing of auditory information that can aid performance on cognitive control tasks. A similar relationship has been suggested in non-human primates where a recent study identified prolonged activity in the auditory cortex to reflect sensorimotor representations important for behavioral inhibition (Huang et al., 2019).

It is noteworthy that even though the right hemisphere showed stronger responses, left hemisphere activity showed the meaningful behavioral association in children. We surmise this relates to the different developmental trajectories of the auditory cortices. The left auditory cortex has been suggested to mature slower than the right (Paetau et al., 1995; Parviainen et al., 2019), based on the later emergence of the N100 response. In addition, auditory responses in the right-hemisphere have been more strongly linked with genetic regulation compared to the left-hemisphere (Renvall et al., 2012). Our reported hemispheric differences may also link to the lateralization of function of the auditory areas, where the left-hemisphere is assumed to have a higher temporal resolution. Finally, handedness has also been shown to affect hemispheric dominance of neuromagnetic responses to sounds (Kirveskari et al., 2006) and as such our reported effect might depend on handedness. An important remaining question is whether our reported relationship depends on the auditory cortex that is contralateral to the hand used to respond, or a mechanism unique to the left hemisphere.

A few theoretical considerations of this study need to be addressed. First, somewhat surprisingly, most brain-behavior correlations did not hold for the passive vs oddball comparison. Arguably, the standard tones used in that comparison were behaviorally less relevant as they never required a response from the participant, which may influence the strength of the brain-behavior relationship. We hypothesize that task relevance of the auditory response is an important factor determining (the strength of) the correlation. This would suggest that the deviant tones are more relevant than the standard tones, perhaps because deviant tones required active inhibition in the context of a Go/No-go task.

Second, we focused our discussion on the ICV and argued it reflects cognitive control. It is, however, good to note that the actual relationship is shown between child specific neural activity in the (left) auditory cortex and individual response time variability in a Go/No-go task. How broadly this can be interpreted, both in terms of other auditory neural responses as well as cognitive processes under the general domain of cognitive control (e.g., selective attention, inhibition or conflict monitoring) is up to debate and should be subject of further investigation.

To conclude, we provide unique evidence that the child-specific auditory activation in the left-hemisphere at around 250ms is functionally meaningful for performance on inhibition tasks. We claim that the mechanisms underlying cognitive control are different in children and adults with more emphasis on sensorimotor associations in children. Interestingly, the correlation between activation strength and performance measures are limited to the left-hemisphere. We presume this reflects the general lateralization of function of the auditory cortices and experience-driven plasticity which is more strongly linked to the left-hemisphere.

## Acknowledgements

We are grateful to Hanna-Maija Lapinkero, Suvi Karjalainen, Maria Vesterinen & Janne Rajaniemi for help with data collection and to Amit Jaiswal, Erkka Heinilä and Jukka Nenonen for their help with preprocessing and scripting. This work was supported by EU project ChildBrain (Horizon2020 Marie Skłodowska-Curie Action (MSCA) Innovative Training Network (ITN) – European Training Network (ETN), grant agreement no. 641652).

